# δ-catenin haploinsufficiency is sufficient to alter behaviors and glutamatergic synapses in mice

**DOI:** 10.1101/2025.08.12.669907

**Authors:** Emma S. Hinchliffe, Victoria Aragon, Van T. Mai, Swapna A. Shah, Rahmi Lee, Jyothi Arikkath, Seonil Kim

## Abstract

δ-catenin (also known as CTNND2) functions as an anchor for the glutamatergic AMPA receptor (AMPARs) to regulate synaptic activity in excitatory synapses. Alteration in the gene coding δ-catenin has been implicated in many neurological disorders. Some of these genetic alterations exhibit a profound loss of δ-catenin functions in excitatory synapses. We have shown that δ-catenin deficiency induced by the homozygous δ-catenin knockout (KO) and autism-associated missense glycine 34 to serine (G34S) mutation significantly alters AMPAR-mediated synaptic activity in cortical neurons and disrupts social behavior in mice. Importantly, many genetic disorders are caused by haploinsufficiency. Indeed, δ-catenin haploinsufficiency contributes to severe autism and learning disabilities in humans. However, previous studies have used only homozygous δ-catenin deficiency models. Therefore, it is important to examine the effects of δ-catenin haploinsufficiency on animals’ behaviors and excitatory synapses. Here, we use heterozygous δ-catenin KO and G34S mice as a δ-catenin haploinsufficiency model to examine this idea. Multiple behavioral assays, a social behavior test, contextual fear conditioning, and an open field test, reveal that both δ-catenin KO and G34S haploinsufficiency significantly disrupt animals’ social behavior and fear learning and memory. Interestingly, only KO haploinsufficiency mice show anxiety-like behavior. A biochemical assay using brain extracts demonstrates that δ-catenin haploinsufficiency significantly affects the levels of synaptic δ-catenin and AMPARs. Our findings thus suggest that δ-catenin haploinsufficiency affects animals’ behaviors via altering glutamatergic synaptic activity.

## Introduction

δ-catenin, also known as CTNND2 or neural plakophilin-related armadillo protein (NPRAP), is a member of the armadillo repeat proteins that is highly expressed in neurons ^1–4^. At the postsynaptic density (PSD), δ-catenin interacts with the intracellular domain of N-cadherin, a synaptic cell adhesion protein, and the carboxyl-terminus of δ-catenin binds to AMPA receptor-binding protein (ABP) and glutamate receptor-interacting protein (GRIP) ^1,4–6^ (**Fig. 1**). This N-cadherin-δ-catenin-ABP/GRIP complex functions as an anchorage for the AMPA-type glutamate receptor (AMPAR) subunit GluA2 ^1^ (**Fig. 1**) and plays a crucial role in maintaining synaptic function and structure ^7^.

**Figure 1.**
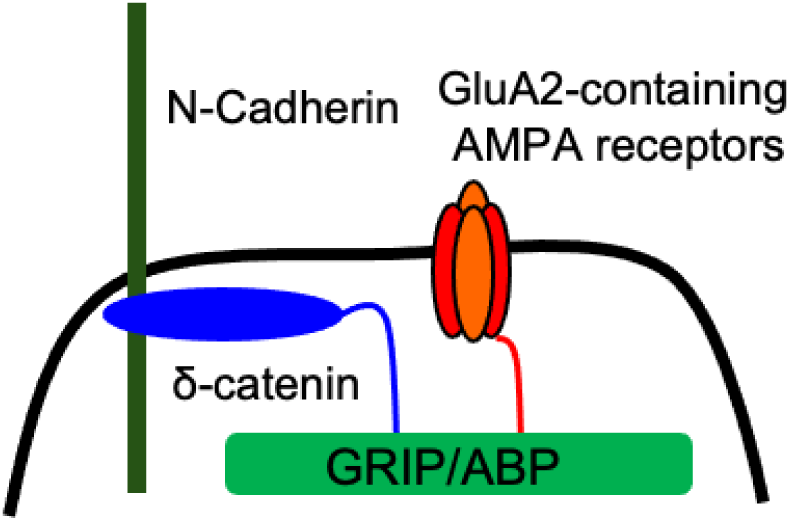
The synaptic N-cadherin-δ-catenin-ABP/GRIP-GluA2 complex. At PSD, δ-catenin interacts with N-cadherin, a synaptic cell adhesion protein. The carboxyl-terminus of δ-catenin binds to AMPA receptor-binding protein (ABP) and glutamate receptor-interacting protein (GRIP). This N-cadherin-δ-catenin-ABP/GRIP complex functions as an anchor for GluA2.

The δ*-catenin* gene in humans harbors deleterious missense mutations, copy number variation, single exon duplication, and de novo mutations that are associated with severely affected autism spectrum disorder (ASD) patients from multiple families ^8–10^. In fact, the δ*-catenin* gene is categorized as a ‘strong candidate’ regarding its association with ASD in the Simons Foundation Autism Research Initiative (SFARI) Database. In addition, the expression of δ-catenin is highly correlated with other ASD-risk genes that are involved in synaptic structure and function, including *cadherins* and *GRIP*, further implicating its potential roles in ASD etiology ^11,12^. Notably, patients heterozygous for deletions or loss-of-function variants of the δ*-catenin* gene exhibit a variety of features of ASD ^8,10,13^. Importantly, we have shown that δ-catenin deficiency induced by δ-catenin knockout (KO) and the ASD-associated missense glycine 34 to serine (G34S) mutation significantly reduces synaptic GluA2 levels in the cortex ^14^. Additionally, we have found altered AMPAR-mediated synaptic activity selectively in cortical neurons ^14^. Recent studies, including our own, suggest that δ-catenin deficiency induced by δ-catenin KO, the G34S mutation, or δ-catenin knockdown significantly reduces cortical neurons’ inhibition to elevate excitation ^14,15^. Research further reveals that the loss of δ-catenin functions disrupts social behavior in mice ^14,16^. Notably, cri du chat syndrome, also known as 5p-syndrome, results from variable hemizygous deletions in the short arm of chromosome 5p, including the δ*-catenin* gene. The clinical features comprise growth delay, severe mental retardation, and speech delay ^17,18^. In those individuals with the syndrome who have mild or no obvious intellectual disability, the deletion often does not include the δ*-catenin* gene ^19^. Moreover, individuals carrying a partial deletion and a partial duplication of the δ*-catenin* gene exhibit cri du chat syndrome, but only mild cognitive disability ^20^. Additionally, a rat model for cri du chat syndrome reveals the important roles of δ-catenin in disease pathogenesis ^21^. This suggests the δ*-catenin* gene plays crucial roles in etiology of the syndrome. Moreover, approximately 40% of individuals with this syndrome show some autistic-like characteristics, including repetitive behavior ^22^. This is considerably higher than expected in the general population ^22^. Importantly, genetic studies identify a loss of δ-catenin function as a significant factor for both cri du chat syndrome and ASD ^10,23–25^. Moreover, δ-catenin mutations have been implicated in many other neurological disorders, including cerebral palsy, schizophrenia, anxiety disorders, attention deficit hyperactivity disorder (ADHD), and Alzheimer’s disease ^26–30^. These findings strongly support the scientific premise that δ-catenin is critical for glutamatergic synaptic activity and pathological behaviors.

Many genetic disorders are caused by haploinsufficiency, in which having only one copy of the wild-type (WT) allele is not sufficient to produce the WT phenotype when the other allele is a loss-of-function mutation or deleted. Indeed, δ-catenin haploinsufficiency contributes to severe ASD and learning disabilities in humans ^8–10,19,31–33^. In previous studies, δ-catenin homozygous KO mice have been used to understand the role of δ-catenin in synaptic plasticity and learning and memory ^8,34^. However, this mouse model contains a truncated form of δ-catenin protein (δ-catenin N-term mice), making it unable to address comprehensive loss-of-function effects. Therefore, we have generated a new δ-catenin KO mouse model that does not contain full-length or truncated δ-catenin ^14^. In fact, our homozygous KO and G34S mice show altered AMPAR-mediated synaptic activity and impaired social behavior ^14^. However, the precise effects of δ-catenin haploinsufficiency on animals’ behaviors and glutamatergic synapses are unknown.

The current study uses heterozygous δ-catenin deficiency mice – KO and G34S – as a haploinsufficiency model. We find that these mice show impairments in social behavior and fear learning and memory. Interestingly, only KO haploinsufficiency mice show anxiety-like behavior. Moreover, we reveal that synaptic δ-catenin and GluA2 levels are significantly reduced in KO haploinsufficiency whole brains when compared to WT and G34S heterozygous mutant brains. Notably, these synaptic changes are also found in the G34S heterozygous mutant cortex. Our findings thus suggest that δ-catenin haploinsufficiency affects animals’ behaviors via altering glutamatergic synaptic activity in various regions of the brain. Therefore, our study suggests that δ-catenin heterozygous mutant is a valuable loss-of-function model to investigate pathophysiology caused by δ-catenin haploinsufficiency.

## Material and Methods

### Animals

δ-catenin KO mouse (C57Bl6J) was developed in collaboration with Cyagen Biosciences Inc. as described previously ^14^. δ-catenin G34S mice (C57Bl6J) (RRID:MMRRC_050621-UCD) were generated by Dr. Jyothi Arikkath at University of Nebraska Medical Center Mouse Genome Engineering Core as described previously ^14^. Both lines were bred and group housed in the animal facility at Colorado State University (CSU). Animals were housed under a 12:12 hour light/dark cycle. All behavior experiments were conducted during the light phase. To collect brain tissues from animals, mice were deeply anesthetized and euthanized by CO_2_ asphyxiation. All efforts were made to minimize animal suffering. CSU’s Institutional Animal Care and Use Committee (IACUC) reviewed and approved the animal care and protocol (6808). Results were reported following the ARRIVE (Animal Research: Reporting of In Vivo Experiments) guidelines ^35^.

### Social behavior

We examined social behavior between two freely moving mice as described previously^36^. A 3-month-old female or male test mouse and a same sex stranger mouse of the identical genotype that was previously housed in different cages were placed into the chamber (40 W x 40 L x 40 H cm) and allowed to explore freely for 20 min. Social behavior was recorded using a camera mounted overhead. Social contacts were determined by the sniffing time that was defined as each instance in which a test mouse’s nose came within 2 cm toward a stranger mouse. The total number and total duration of social contacts were measured manually. Animals that escaped the chamber or showed aggressive behaviors (e.g. attacking) were excluded from the analysis.

### Open field test (OFT)

We measured locomotor activity and anxiety-like behavior using the OFT as carried out previously ^36–38^. A 3-month-old female or male test mouse was first habituated in the open field chamber (40 W x 40 L x 40 H cm) for 5 min. Animals were then allowed to explore the chamber for additional 20 min. The behavior was recorded by a video camera mounted overhead. A 20 x 20 cm center square was defined as the inside zone. Data were analyzed using the ANY-maze tracking program (Stoelting Co.) to acquire the total traveled distance (locomotor activity) and the time spent outside and inside (anxiety-like behavior). Animals that escaped the chamber or showed significantly decreased locomotor activity were excluded from the analysis.

### Contextual fear conditioning (CFC)

CFC (Habitest Modular System, Coulbourn Instrument) was carried out as described previously ^36,37,39,40^. On Day 1, a 3-month-old female or male test mouse was placed in a novel rectangular chamber with a grid floor. After a 3-min baseline period, the test animal was given one shock (a 2 sec, 0.5 mA shock) and stayed in the chamber for an additional 1 min after the shock before being returned to the home cage. A contextual memory test was conducted the next day (Day 2) in the same conditioning chamber for 3 min. Fear memory was determined by measuring the percentage of the freezing response (immobility excluding respiration and heartbeat) using an automated tracking program (FreezeFrame).

### Tissue sample preparation and immunoblots

We used whole brains or the cortex from 3-month-old female and male mice and isolated whole tissue lysates and the PSD fraction as shown previously ^14,41^. Brain tissues were rinsed with PBS, collected to 15 ml tubes, homogenized by douncing in solution A (320 mM Sucrose, 1 mM NaHCO_3_ 1 mM MgCl_2_, 0.5 mM CaCl_2_, 0.1 mM PMSF, and 1X protease inhibitor cocktail), and spun at 2,000 rpm for 10 min at 4°C. The supernatant (*whole tissue lysates*) was centrifuged at 13,000 rpm for 20 min at 4°C. The pellet was homogenized in solution B (320 mM Sucrose and 1 mM NaHCO_3_), placed on top of a 1 M sucrose and 1.2 M sucrose gradient, and centrifuged at 28,000 rpm for 2 hours at 4°C. The resulting interface between the gradient was centrifuged at 40,000 rpm for 45 min at 4°C after a six-fold dilution with solution B. The pellet was resuspended in 1% Triton-X and 25 mM Tris buffer, inculcated for 30 min at 4°C while rocking, and spun at 13,000 rpm for 30 min at 4°C. The pellet (*PSD*) was resuspended in 2% SDS and 25 mM Tris buffer. The protein concentration in each faction was determined by NanoDrop Lite (Thermo Fisher Scientific). Immunoblots were performed as described previously ^14,37,39,41–54^. Equal amounts of protein samples were loaded on 10% glycine-SDS-PAGE gel. The gels were transferred to nitrocellulose membranes. The membranes were blocked (5% powdered milk) for 1 hour at room temperature, followed by overnight incubation with the primary antibodies at 4°C. The primary antibodies consisted of anti-δ-catenin (BD Biosciences, 1:1000, 611537), anti-GluA1 (RH95) (Millipore, 1:2000, MAB2263), anti-GluA2 (EPR18115) (Abcam, 1:2000, ab206293), and anti-actin (Abcam, 1:2000, ab3280) antibodies. The membrane was developed with chemiluminescence (Life Tech, PI 34580). Protein bands were quantified using the NIH ImageJ software (https://imagej.nih.gov/ij/). We repeated 4 times with the same samples for statistical analysis.

### Statistical analysis

All behavior tests were blindly scored by more than two investigators. The Franklin A. Graybill Statistical Laboratory at Colorado State University was consulted for statistical analysis in the current study, including sample size determination, randomization, experiment conception and design, data analysis, and interpretation. We used the GraphPad Prism 10 software to determine statistical significance (set at *p* < 0.05). Grouped results of single comparisons will be tested for normality with the Shapiro-Wilk normality or Kolmogorov-Smirnov test and analyzed using a two-tailed Student’s t-test when data are normally distributed. Differences between three groups were assessed by One-way analysis of variance (ANOVA) with the Tukey test. The graphs were presented as mean ± Standard Deviation (SD). We discard data points that are located further than two SD above or below the average as an outlier.

## Results

### δ-catenin haploinsufficiency is sufficient to disrupt social behavior in mice

We have shown that δ-catenin deficiency induced by the homozygous knockout (KO) and the ASD-associated missense G34S mutation disrupts social behavior ^14^. We thus examined whether δ-catenin haploinsufficiency disrupted social behavior between two freely moving mice by comparing social contacts between WT, heterozygous KO (KO HET), and heterozygous G34S (G34S HET) mice. We found that δ-catenin haploinsufficiency significantly decreased the total number of social contacts (FEMALE: WT, 50.80 ± 25.72, KO HET, 33.18 ± 17.67, and G34S HET, 24.85 ± 10.65, WT vs. KO HET: *p* = 0.0086 and WT vs. G34S HET: *p* < 0.0001. MALE: WT, 57.06 ± 37.72, KO HET, 32.00 ± 15.31, and G34S HET, 34.71 ± 18.47, WT vs. KO HET: *p* = 0.0273 and WT vs. G34S HET: *p* = 0.0147) (**Fig. 2 and Supplementary Table 1**) and total time of social contacts (FEMALE: WT, 60.01 ± 40.64 sec, KO HET, 37.06 ± 27.49 sec, and G34S HET, 23.36 ± 13.05 sec, WT vs. KO HET: *p* = 0.0303 and WT vs. G34S HET: *p* < 0.0001. MALE: WT, 60.86 ± 35.87 sec, KO HET, 37.92 ± 21.08 sec, and G34S HET, 35.75 ± 20.31 sec, WT vs. KO HET: *p* = 0.0468 and WT vs. G34S HET: *p* = 0.0052) (**Fig. 2 and Supplementary Table 2**) in both female and male δ-catenin haploinsufficiency mice, an indication of social dysfunction. These findings demonstrate that δ-catenin haploinsufficiency is sufficient to impair animals’ social behavior.

**Figure 2.**
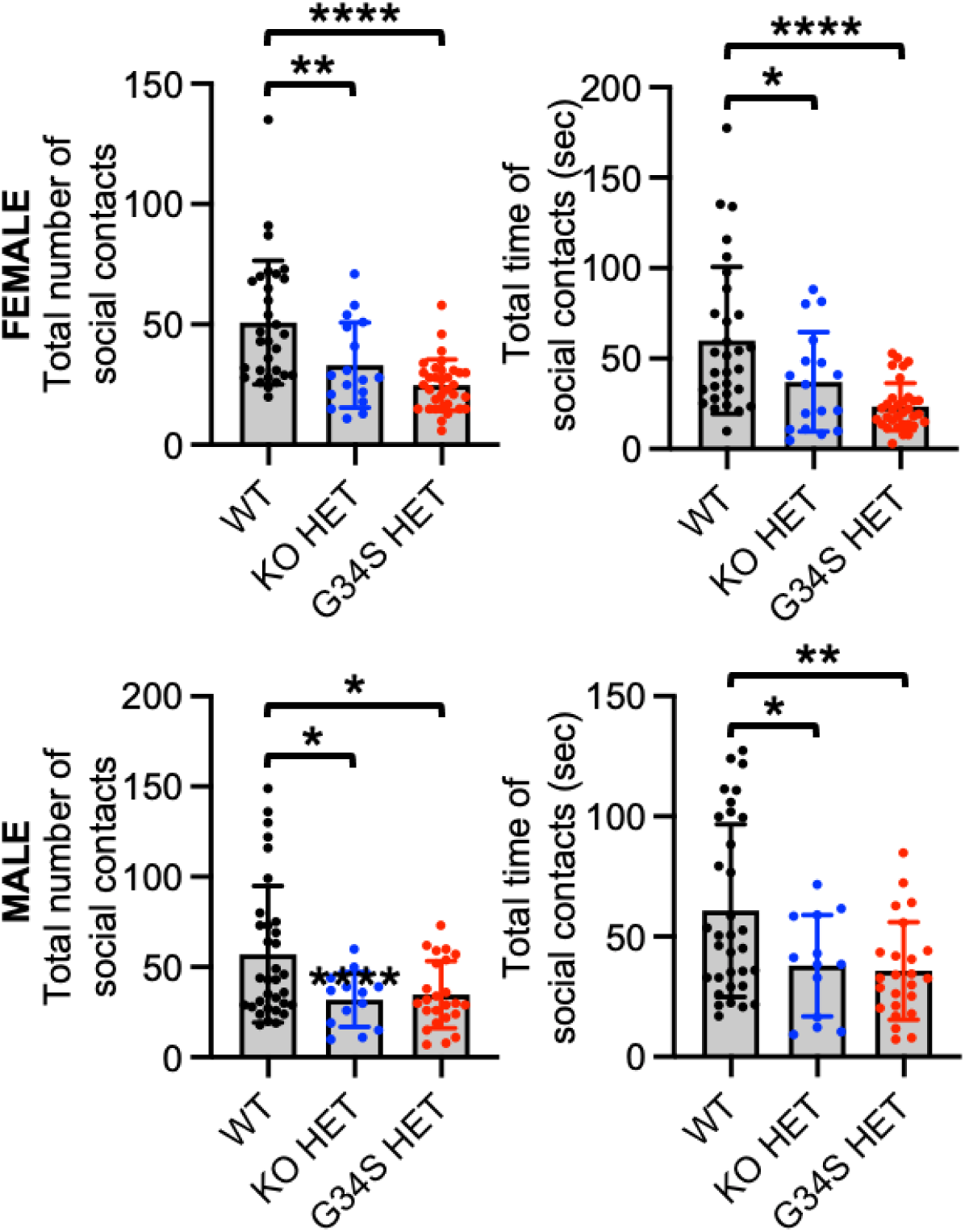
δ-catenin haploinsufficiency is sufficient to disrupt social behavior in mice. Summary of total numbers and time of social contacts in each condition (n = number of animals. FEMALE: WT = 30, KO HET = 17, and G34S HET = 33. MALE: WT = 33, KO HET = 13, and G34S HET = 24). \**p* < 0.05, \*\**p* < 0.01, and \*\*\*\**p* < 0.0001, One-Way ANOVA, Tukey test.

### Heterozygous **δ**-catenin KO mice exhibit reduced locomotor activity and anxiety-like behavior

We next examined whether δ-catenin haploinsufficiency affects locomotor activity anxiety-like behavior in mice using the OFT ^14,36–38^. We measured the total distance traveled (locomotor activity) and the total time spent outside and inside (anxiety-like behavior) in the open field chamber. Interestingly, KO HET significantly decreased the total distance travelled compared to WT and G34S HET in both female and male mice, an indication of hypolocomotion (FEMALE: WT, 40.50 ± 15.01 m, KO HET, 28.97 ± 16.69 m, and G34S HET, 46.86 ± 16.26 m. WT vs. KO HET: *p* = 0.0155 and KO HET vs. G34S HET: *p* = 0.0002. MALE: WT, 38.41 ± 13.89 m, KO HET, 23.66 ± 12.96 m, and G34S HET, 33.75 ± 11.39 m. WT vs. KO HET: *p* = 0.0001 and KO HET vs G34S HET: *p* = 0.0257) (**Fig. 3 and Supplementary Table 3**). Furthermore, KO HET mice spent more time outside (FEMALE: WT, 1022 ± 123.0 sec, KO HET, 1136 ± 33.68 sec, and G34S HET, 1024 ± 78.26 sec. WT vs. KO HET: *p* < 0.0001 and KO HET vs. G34S HET: *p* < 0.0001. MALE: WT, 1055 ± 99.64 sec, KO HET, 1140 ± 12.96 sec, and G34S HET, 1052 ± 68.25 sec. WT vs. KO HET: *p* = 0.0003 and KO HET vs G34S HET: *p* = 0.0008) (**Fig. 3 and Supplementary Table 4**) but less time inside (FEMALE: WT, 174.4 ± 123.3 sec, KO HET, 62.16 ± 33.78 sec, and G34S HET, 175.6 ± 78.26 sec. WT vs. KO HET: *p* < 0.0001 and KO HET vs. G34S HET: *p* < 0.0001. MALE: WT, 145.4 ± 99.64 sec, KO HET, 59.51 ± 33.03 sec, and G34S HET, 147.7 ± 68.25 sec. WT vs. KO HET: *p* = 0.0003 and KO HET vs G34S HET: *p* = 0.0008) than WT and G34S HET mice (**Fig. 3 and Supplementary Table 5**), indicating anxiety-like behavior. This indicates that KO HET in mice significantly decreases locomotor activity and induces anxiety-like behavior in the OFT while G34S HET has no effect on these behaviors.

**Figure 3.**
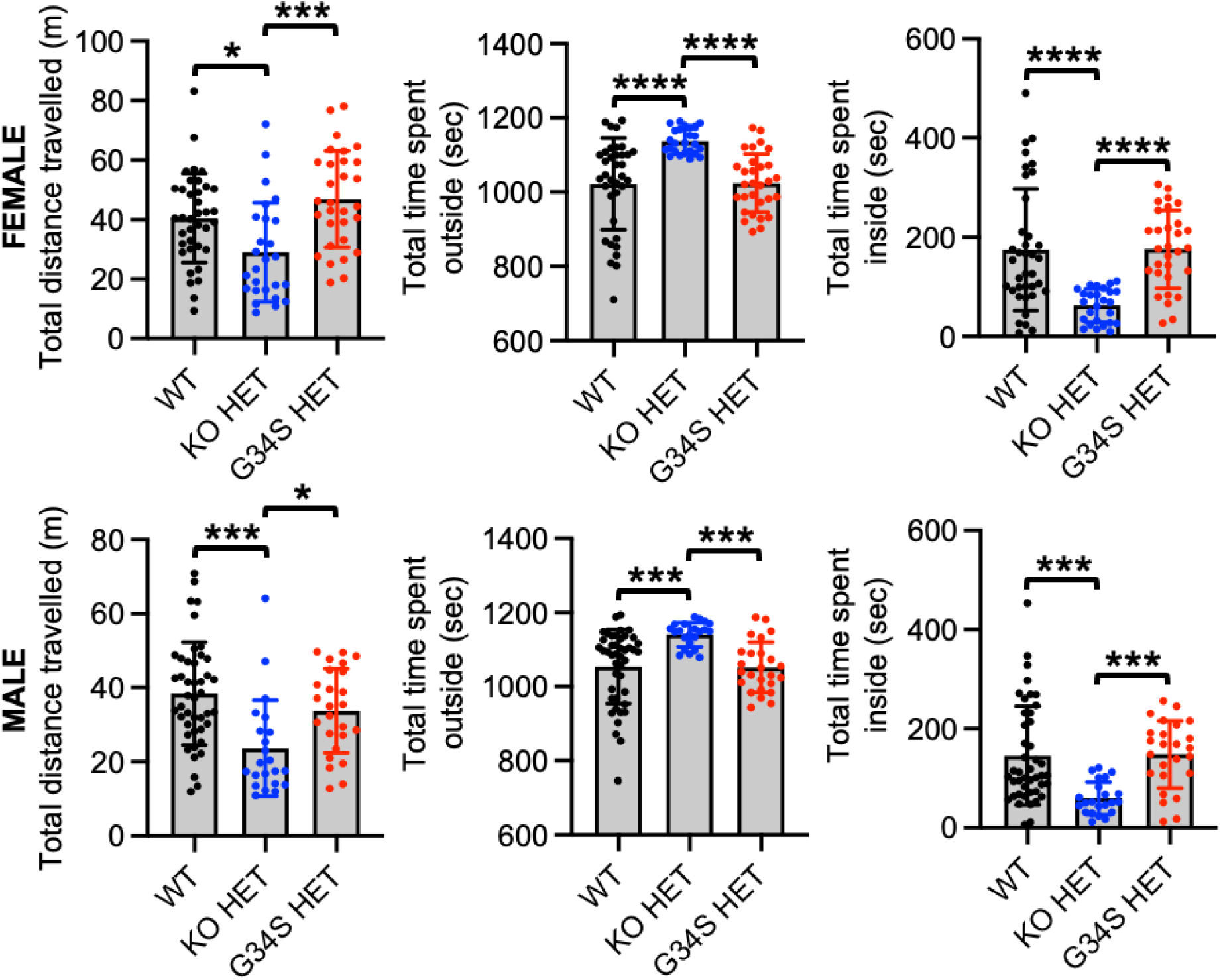
Heterozygous δ-catenin KO mice exhibit reduced locomotor activity and anxiety-like behavior. Summary of total distance traveled, total time spent outside, and total time spent inside in each condition (n = number of animals. FEMALE: WT = 37, KO HET = 26, and G34S HET = 30. MALE: WT = 44, KO HET = 22, and G34S HET = 25). \**p* < 0.05, \*\*\**p* < 0.001, and \*\*\*\**p* < 0.0001, One-Way ANOVA, Tukey test.

### **δ**-catenin haploinsufficiency is sufficient to disrupt fear learning and memory

A loss of δ-catenin is shown to disrupt spatial learning and memory in animals ^16,21^. Contextual fear learning and memory is also impaired in δ-catenin N-term mice ^34^. However, the role of δ-catenin haploinsufficiency in learning and memory has not been determined. We thus used CFC to examine whether δ-catenin haploinsufficiency disrupted hippocampus-dependent fear learning and memory in mice. We found that freezing was significantly lower in both female and male δ-catenin haploinsufficiency mice compared to WT controls, an indication of fear learning and memory loss (FEMALE: WT, 34.00 ± 14.13%, KO HET, 21.66 ± 10.34%, and G34S HET, 22.08 ± 8.141%, WT vs. KO HET: *p* = 0.0211 and WT vs. G34S HET: *p* = 0.0452. MALE: WT, 38.77 ± 10.56%, KO HET, 25.26 ± 9.640%, and G34S HET, 22.19 ± 13.16%, WT vs. KO HET: *p* = 0.0174 and WT vs. G34S HET: *p* = 0.0023) (**Fig. 4 and Supplementary Table 6**). This data suggest that δ-catenin haploinsufficiency disrupts hippocampus-dependent fear learning and memory in animals.

**Figure 4.**
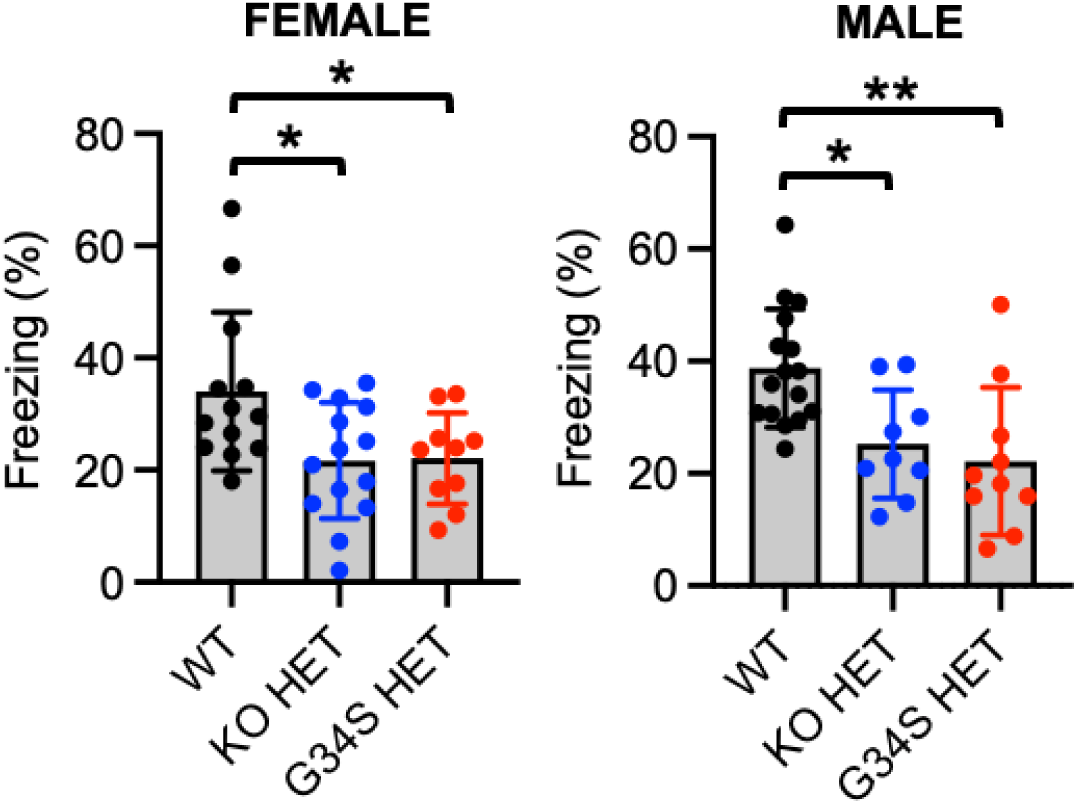
δ-catenin haploinsufficiency is sufficient to disrupt fear learning and memory. Summary of fear memory in each condition (n = number of animals. FEMALE: WT = 13, KO HET = 14, and G34S HET = 10. MALE: WT = 16, KO HET = 9, and G34S HET = 10). \**p* < 0.05 and \*\**p* < 0.01, One-Way ANOVA, Tukey test.

### Synaptic **δ**-catenin and GluA2 levels are significantly reduced in heterozygous **δ**-catenin KO brains

We have previously shown that synaptic δ-catenin and GluA2 levels are significantly decreased in the homozygous δ-catenin KO and G34S cortex while there is no difference in these proteins in cortical whole tissue lysates ^14^. Given that AMPAR subunit composition and trafficking allow for complex regulation of synaptic strength and ultimately impacts various behavioral outputs in animals ^55^, we examined whether δ-catenin haploinsufficiency affected δ-catenin and AMPAR levels in whole brains. The whole tissue lysates and PSD fractions of whole brains were collected from 3-month-old female and male WT and δ-catenin haploinsufficiency animals and analyzed for total and synaptic protein levels, respectively, using immunoblotting. For whole tissue lysates, we found significant reduction in δ-catenin levels in female and male KO HET whole brains when compared to WT and G34S HET tissues (FEMALE: WT, 1.000 ± 0.378, KO HET, 0.493 ± 0.172, and G34S HET, 1.160 ± 0.491. WT vs. KO HET: *p* = 0.0055 and KO HET vs. G34S HET: *p* = 0.0003. MALE: WT, 1.000 ± 0.425, KO HET, 0.685 ± 0.257, and G34S HET, 0.990 ± 0.167. WT vs. KO HET: *p* = 0.0400 and KO HET vs. G34S HET: *p* = 0.0481) (**Fig. 5a and 5b and Supplementary Table 7**). However, no significant difference was found in GluA1 (FEMALE: WT, 1.000 ± 0.415, KO HET, 0.730 ± 0.342, and G34S HET, 0.741 ± 0.549. MALE: WT, 1.000 ± 0.456, KO HET, 0.914 ± 0.434, and G34S HET, 1.137 ± 0.461) and GluA2 levels (FEMALE: WT, 1.000 ± 0.460, KO HET, 0.900 ± 0.504, and G34S HET, 1.216 ± 0.863. MALE: WT, 1.000 ± 0.366, KO HET, 1.154 ± 0.361, and G34S HET, 1.207 ± 0.476) between the groups (**Fig. 5a and 5b and Supplementary Table 7**). These findings show that total δ-catenin levels are significantly reduced only in KO HET brains while there is no difference in total GluA1 and GluA2 levels throughout the groups. Next, we found a significant reduction in synaptic δ-catenin only in female KO HET brains when compared to WT and G34S HET female tissues (FEMALE: WT, 1.000 ± 0.221, KO HET, 0.380 ± 0.161, and G34S HET, 0.822 ± 0.372. WT vs. KO HET: *p* < 0.0001 and KO HET vs. G34S HET: *p* = 0.0008) (**Fig. 5c and 5d and Supplementary Table 8**). Interestingly, synaptic δ-catenin levels in both KO and G34S HET male brains were markedly decreased when compared to WT controls (MALE: WT, 1.000 ± 0.356, KO HET, 0.680 ± 0.219, and G34S HET, 0.673 ± 0.267. WT vs. KO HET: *p* = 0.0267 and WT vs. G34S HET: *p* = 0.0231), suggesting the sex difference effects on synaptic δ-catenin levels in δ-catenin haploinsufficiency brains (**Fig. 5c and 5d and Supplementary Table 8**). We also revealed that synaptic GluA2 levels were significantly reduced only in KO HET brains compared to WT and G34S HET groups (FEMALE: WT, 1.000 ± 0.190, KO HET, 0.671 ± 0.163, and G34S HET, 0.973 ± 0.359. WT vs. KO HET: *p* = 0.0084 and KO HET vs. G34S HET: *p* = 0.0164. MALE: WT, 1.000 ± 0.151, KO HET, 0.693 ± 0.206, and G34S HET, 1.180 ± 0.436. WT vs. KO HET: *p* = 0.0382 and KO HET vs. G34S HET: *p* = 0.0008) (**Fig. 5c and 5d and Supplementary Table 8**). Interestingly, we discovered that there was no difference in synaptic GluA1 levels between WT and KO HET brains, while synaptic GluA1 in G34S HET brains was significantly elevated when compared to other groups (FEMALE: WT, 1.000 ± 0.125, KO HET, 0.820 ± 0.275, and G34S HET, 1.615 ± 0.900. WT vs. G34S HET: *p* = 0.0253 and KO HET vs. G34S HET: *p* = 0.0032. MALE: WT, 1.000 ± 0.222, KO HET, 0.754 ± 0.276, and G34S HET, 1.557 ± 0.834. WT vs. G34S HET: *p* = 0.0354 and KO HET vs. G34S HET: *p* = 0.0019) (**Fig. 5c and 5d and Supplementary Table 8**). These findings suggest that synaptic δ-catenin and GluA2 levels are significantly reduced in addition to total δ-catenin levels in KO HET whole brains, an indication of AMPAR-mediated synaptic dysfunction. Conversely, G34S HET whole brains show some moderate changes of δ-catenin and AMPAR levels only in synapses.

**Figure 5.**
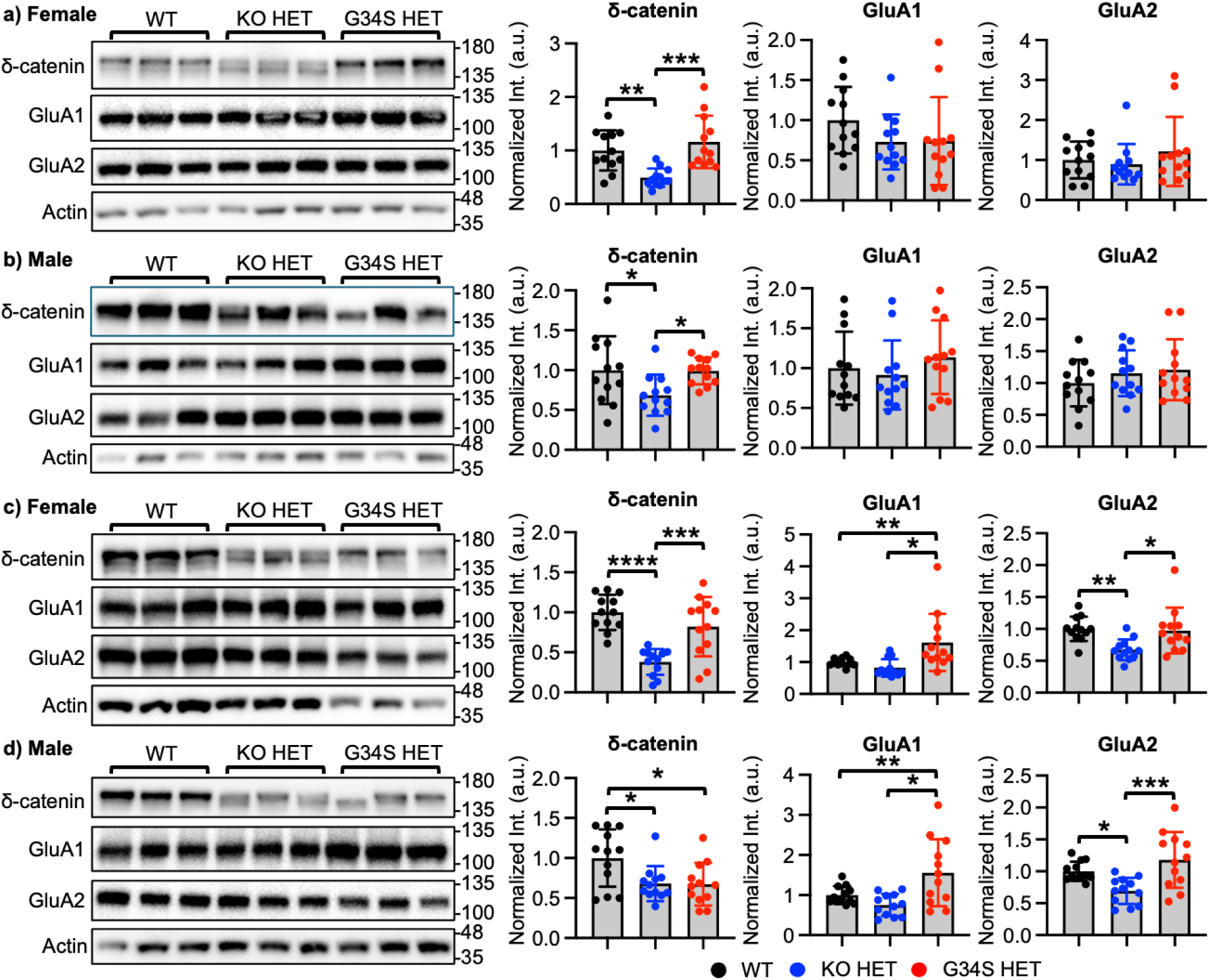
Synaptic δ-catenin and GluA2 levels are significantly reduced in heterozygous δ-catenin KO brains. **a-b)** Representative immunoblots of total δ-catenin and AMPAR levels and summary graphs of normalized δ-catenin, GluA1, and GluA2 in each condition in whole brains. **c-d)** Representative immunoblots of synaptic δ-catenin and AMPAR levels and summary graphs of normalized δ-catenin, GluA1, and GluA2 in each condition in the PSD fraction. n = 12 immunoblots from 3 mice in each condition. \**p* < 0.05, \*\**p* < 0.01, \*\*\**p* < 0.001, and \*\*\*\**p* < 0.0001, One-Way ANOVA, Tukey test. The position of molecular mass markers (kDa) is shown on the right of the blots.

### Synaptic **δ**-catenin and GluA2 levels are significantly reduced in the heterozygous **δ**-catenin G34S cortex

Although we found significant differences in G34S HET animals’ behaviors (**Fig. 2 and 4**), there were only moderate changes in synaptic δ-catenin and AMPAR levels in whole brains unlike KO HET brains (**Fig. 5**). Given that these proteins are significantly decreased in the homozygous G34S cortex ^14^, we isolated the cortex from WT and G34S HET mice and examined whether δ-catenin G34S haploinsufficiency affected δ-catenin and AMPAR levels specifically in the cortex. Similar to the previous findings in the homozygous G34S cortex ^14^, total δ-catenin (FEMALE: WT, 1.000 ± 0.146 and G34S HET, 1.220 ± 0.424, t = 1.473, df = 16, *p* = 0.1602. MALE: WT, 1.000 ± 0.423 and G34S HET, 0.800 ± 0.285, t = 1.189, df = 16, *p* = 0.2516), GluA1 (FEMALE: WT, 1.000 ± 0.194 and G34S HET, 1.122 ± 0.539, t = 0.641, df = 16, *p* = 0.5304. MALE: WT, 1.000 ± 0.201 and G34S HET, 1.366 ± 0.505, t = 2.021, df = 16, *p* = 0.0604), and GluA2 levels (FEMALE: WT, 1.000 ± 0.171 and G34S HET, 0.870 ± 0.353, t = 1.000, df = 16, *p* = 0.3324. MALE: WT, 1.000 ± 0.239 and G34S HET, 1.174 ± 0.647, t = 0.757, df = 16, *p* = 0.4599) were unaltered in the G34S HET cortex when compared to WT controls (**Fig. 6a and 6b**). However, the G34S HET cortex exhibited a significant reduction in synaptic δ-catenin (FEMALE: WT, 1.000 ± 0.387 and G34S HET, 0.393 ± 0.188, t = 4.231, df = 16, *p* = 0.0006. MALE: WT, 1.000 ± 0.327 and G34S HET, 0.281 ± 0.183, t = 5.759, df = 16, *p* < 0.0001), GluA1 (FEMALE: WT, 1.000 ± 0.438 and G34S HET, 0.462 ± 0.130, t = 3.533, df = 16, *p* = 0.0028. MALE: WT, 1.000 ± 0.170 and G34S HET, 0.722 ± 0.301, t = 2.412, df = 16, *p* = 0.0283), and GluA2 levels (FEMALE: WT, 1.000 ± 0.161 and G34S HET, 0.615 ± 0.259, t = 3.797, df = 16, *p* = 0.0016. MALE: WT, 1.000 ± 0.265 and G34S HET, 0.649 ± 0.386, t = 2.257, df = 16, *p* = 0.0383) compared to these protein levels in the WT cortex (**Fig. 6c and 6d**). These findings suggest that synaptic δ-catenin and GluA2 levels are significantly reduced in cortical G34S HET tissues, an indication of AMPAR-mediated synaptic dysfunction specifically in the cortex.

**Figure 6.**
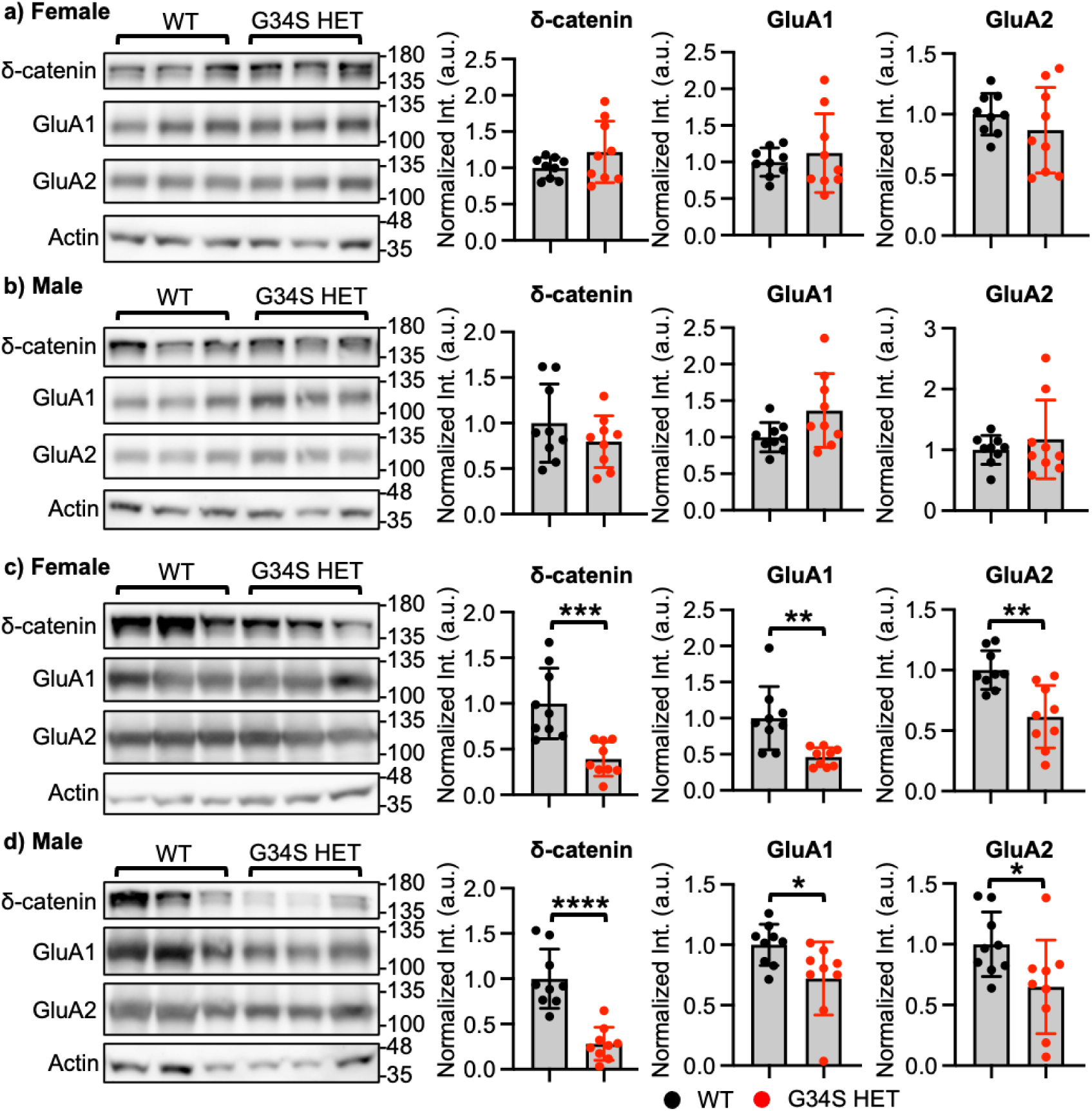
Synaptic δ-catenin and GluA2 levels are significantly reduced in the heterozygous δ-catenin G34S cortex. **a-b)** Representative immunoblots of total δ-catenin and AMPAR levels and summary graphs of normalized δ-catenin, GluA1, and GluA2 in each condition in the cortex. **c-d)** Representative immunoblots of synaptic δ-catenin and AMPAR levels and summary graphs of normalized δ-catenin, GluA1, and GluA2 in each condition in the cortical PSD fraction. n = 12 immunoblots from 3 mice in each condition. \**p* < 0.05, \*\**p* < 0.01, \*\*\**p* < 0.001, and \*\*\*\**p* < 0.0001, two-tailed Student’s t-test. The position of molecular mass markers (kDa) is shown on the right of the blots.

## Discussion

δ-catenin haploinsufficiency is associated with several neurodevelopmental and neuropsychiatric disorders such as intellectual disability, ASD, cri du chat syndrome, schizophrenia, and ADHD ^26–30^. Although several δ-catenin deficiency animal models have been developed to investigate the role of δ-catenin in these pathological conditions ^14–16,21,34^, the effects of δ-catenin haploinsufficiency have not been investigated. Here, we employ the heterozygous KO and ASD-associated G34S mutation as a δ-catenin haploinsufficiency model to demonstrate that these δ-catenin haploinsufficiency models are sufficient to disrupt social contacts and fear learning and memory. Additionally, KO HET mice exhibit hypolocomotion and anxiety-like behavior. Interestingly, synaptic δ-catenin and GluA2 levels are significantly reduced in KO HET brains while these proteins are markedly decreased in the G34S HET cortex. These results suggest that glutamatergic synaptic activity in different parts of the brain is altered by δ-catenin haploinsufficiency, which affects behaviors in mice. Furthermore, our findings indicate that a heterozygous δ-catenin mutant (KO and G34S) is a useful loss-of-function model for examining the pathophysiology brought on by δ-catenin haploinsufficiency.

ASD often involves differences in social behavior, but the exact patterns can vary widely between individuals ^56^. These differences are linked to both neurological and developmental factors, and they can influence how people with ASD perceive, interpret, and respond to social cues ^56^. In ASD animal models, social contacts are measured to evaluate ASD-related social behavior differences ^57^. Since several genes linked to ASD and environmental factors can affect how animals initiate and maintain social interaction, this is a crucial behavioral domain in preclinical ASD research. ^57^. The δ*-catenin* gene in humans has harmful missense mutations, copy number variations, single exon duplications, and de novo mutations that are linked to individuals with severe ASD ^8–10^. Notably, individuals who are heterozygous for δ-catenin gene deletions or loss-of-function variations have a range of ASD symptoms ^8,10,13^. Importantly, our previous report has shown that δ-catenin deficiency induced by homozygous δ-catenin KO and ASD-associated G34S mutation disrupts social behavior ^14^. In the current study, we further demonstrate that δ-catenin haploinsufficiency is sufficient to impair social contacts in KO and G34S HET animals (**Fig. 2**). These findings demonstrate crucial roles of δ-catenin in social behavior.

Studies estimate that 40–50% of ASD children and adults meet diagnostic criteria for at least one anxiety disorder ^58^. Anxiety in ASD can worsen social difficulties (e.g., avoidance leads to fewer practice opportunities) and can amplify repetitive behaviors ^59^. Importantly, several mouse models of ASD exhibit anxiety-like behaviors, often characterized by increased self-grooming and altered responses in tests like the OFT and elevated plus maze ^60^. Importantly, we discover that KO HET mice exhibit anxiety-like behavior (**Fig. 3**), suggesting that KO-mediated δ-catenin haploinsufficiency is sufficient to cause anxiety-like behavior. Conversely, G34S HET mice show normal behavior in the OFT (**Fig. 3**). This suggests that G34S and δ-catenin KO have differential effects on anxiety-like behavior.

Evidence from genetic studies in humans and animal models strongly supports a link between the δ*-catenin* gene and intellectual disability ^19^. Understanding the precise mechanisms involved can potentially lead to new avenues for diagnosis and treatment for individuals affected by δ-catenin-related intellectual disability. In fact, previous reports using various δ-catenin deficiency animal models have shown impairments in learning and memory ^16,21,34^. However, the role of δ-catenin haploinsufficiency in cognitive function has not been previously determined. Our current study using KO and G34S HET mice reveals that δ-catenin haploinsufficiency is sufficient to disrupt contextual fear learning and memory (**Fig. 4**).

This δ-catenin complex functions as an anchorage for GluA2 ^1^ (**Fig. 1**), which is important for synaptic function and structure ^7^. Our previous study has showed that accelerated degradation of synaptic δ-catenin in the cortex caused by the homozygous G34S mutation linked to ASD results in δ-catenin deficiency ^14^. The current work reveals that total δ-catenin levels are significantly reduced only in the KO HET brains when compared to WT and G34S HET tissues (**Fig. 5a and 5b**). Although total δ-catenin levels are significantly reduced in KO HET brains, total GluA1 and GluA2 levels are not altered in both KO and G34S HET tissues compared to WT controls (**Fig. 5a and 5b**), suggesting that δ-catenin haploinsufficiency is unable to affect total AMPAR levels. However, in the PSD fraction from whole KO HET brains, synaptic δ-catenin and GluA2 levels are markedly reduced when compared to WT and G34S HET synapses (**Fig. 5c and 5d**). Notably, in the PSD fraction from whole G34S HET brains, synaptic δ-catenin levels are significantly reduced only in male mice compared to WT controls while they are unaltered in synapses from female whole brains (**Fig. 5c and 5d**). This sex difference has also been found in the cortex of homozygous G34S mice ^14^. This suggests that animal sex may differentially control δ-catenin levels, which is a crucial area for further study. Unexpectedly, we discover that synaptic GluA1 levels are significantly elevated in G34S HET brains when compared to WT and KO HET synapses (**Fig. 5c and 5d**). This change can be mediated by developmental compensation of a G34S HET-induced loss of δ-catenin functions in brains. Although no alteration of synaptic AMPARs in G34S HET whole brains is found, G34S HET animals exhibit behavioral alterations. Given that synaptic δ-catenin and GluA2 levels are significantly decreased in the homozygous G34S cortex ^14^, we examine the G34S HET cortex and discover that synaptic δ-catenin, GluA1, and GluA2 levels are markedly reduced when compared to WT controls while total protein levels are unaltered (**Fig. 6**). According to these findings, KO HET has effects on δ-catenin and its related AMPARs in whole brains, while G34S HET probably affects these proteins specifically in cortical synapses. This likely causes differential effects of G34S and δ-catenin KO on anxiety-like behavior.

In conclusion, according to our findings, we suggest that animals with δ-catenin haploinsufficiency exhibit pathological behaviors because of changes in glutamatergic synaptic activity in different parts of the brain. Therefore, our study shows that a heterozygous δ-catenin mutant is a useful loss-of-function model for examining the pathophysiology brought on by δ-catenin haploinsufficiency.

## Acknowledgements

We thank members of the Kim laboratory for their generous support. This work is supported by NIH T34GM140958, NIH R01MH132921, and the Boettcher foundation.

