## Supplementary Table for "δ-catenin haploinsufficiency is sufficient to alter behaviors and glutamatergic synapses in mice"

**Supplementary Table 1. Statistical analysis of the total number of social contacts.**

| **Female** |  | | | | |
| --- | --- | --- | --- | --- | --- |
| ANOVA summary |  |  | | | |
| F | 14.96 |  |  |  |  |
| P value | <0.0001 |  |  |  |  |
| P value summary | **** |  |  |  |  |
| Significant diff. among means (P < 0.05)? | Yes |  |  |  |  |
| Tukey's multiple comparisons test | Mean diff. | | 95.00% CI of diff. | Adjusted *P* Value | |
| WT vs. KO HET | 17.62 | | 3.837 to 31.41 | 0.0086 | |
| WT vs. G34S HET | 25.95 | | 14.50 to 37.41 | <0.0001 | |
| KO HET vs. G34S HET | 8.328 | | -5.230 to 21.89 | 0.3119 | |
| **Male** |  | | | | |
| ANOVA summary |  |  | | | |
| F | 5.684 |  |  |  |  |
| P value | 0.0052 |  |  |  |  |
| P value summary | ** |  |  |  |  |
| Significant diff. among means (P < 0.05)? | Yes |  |  |  |  |
| Tukey's multiple comparisons test | Mean diff. | | 95.00% CI of diff. | | Adjusted *P* Value |
| WT vs. KO HET | 25.06 | | 2.331 to 47.79 | | 0.0273 |
| WT vs. G34S HET | 22.35 | | 3.731 to 40.97 | | 0.0147 |
| KO HET vs. G34S HET | -2.708 | | -26.61 to 21.20 | | 0.9602 |

**Supplementary Table 2. Statistical analysis of the total time of social contacts.**

| **Female** |  | | | | |
| --- | --- | --- | --- | --- | --- |
| ANOVA summary |  |  | | | |
| F | 12.53 |  |  |  |  |
| P value | <0.0001 |  |  |  |  |
| P value summary | **** |  |  |  |  |
| Significant diff. among means (P < 0.05)? | Yes |  |  |  |  |
| Tukey's multiple comparisons test | Mean diff. | | 95.00% CI of diff. | Adjusted *P* Value | |
| WT vs. KO HET | 22.94 | | 1.793 to 44.09 | 0.0303 | |
| WT vs. G34S HET | 36.64 | | 19.07 to 54.22 | <0.0001 | |
| KO HET vs. G34S HET | 13.70 | | -7.098 to 34.50 | 0.2629 | |
| **Male** |  | | | | |
| ANOVA summary |  |  | | | |
| F | 5.684 |  |  |  |  |
| P value | 0.0052 |  |  |  |  |
| P value summary | ** |  |  |  |  |
| Significant diff. among means (P < 0.05)? | Yes |  |  |  |  |
| Tukey's multiple comparisons test | Mean diff. | | 95.00% CI of diff. | | Adjusted *P* Value |
| WT vs. KO HET | 25.06 | | 2.331 to 47.79 | | 0.0273 |
| WT vs. G34S HET | 22.35 | | 3.731 to 40.97 | | 0.0147 |
| KO HET vs. G34S HET | -2.708 | | -26.61 to 21.20 | | 0.9602 |

**Supplementary Table 3. Statistical analysis of the total distance travelled.**

| **Female** |  | | | | |
| --- | --- | --- | --- | --- | --- |
| ANOVA summary |  |  | | | |
| F | 8.992 |  |  |  |  |
| P value | 0.0003 |  |  |  |  |
| P value summary | *** |  |  |  |  |
| Significant diff. among means (P < 0.05)? | Yes |  |  |  |  |
| Tukey's multiple comparisons test | Mean diff. | | 95.00% CI of diff. | Adjusted *P* Value | |
| WT vs. KO HET | 11.53 | | 1.840 to 21.23 | 0.0155 | |
| WT vs. G34S HET | -6.360 | | -15.67 to 2.948 | 0.2390 | |
| KO HET vs. G34S HET | -17.89 | | -28.05 to -7.744 | 0.0002 | |
| **Male** |  | | | | |
| ANOVA summary |  |  | | | |
| F | 9.404 |  |  |  |  |
| P value | 0.0002 |  |  |  |  |
| P value summary | *** |  |  |  |  |
| Significant diff. among means (P < 0.05)? | Yes |  |  |  |  |
| Tukey's multiple comparisons test | Mean diff. | | 95.00% CI of diff. | | Adjusted *P* Value |
| WT vs. KO HET | 14.75 | | 6.641 to 22.86 | | 0.0001 |
| WT vs. G34S HET | 4.661 | | -3.118 to 12.44 | | 0.3308 |
| KO HET vs. G34S HET | -10.09 | | -19.17 to -1.010 | | 0.0257 |

**Supplementary Table 4. Statistical analysis of the time spent outside.**

| **Female** |  | | | | |
| --- | --- | --- | --- | --- | --- |
| ANOVA summary |  |  | | | |
| F | 13.96 |  |  |  |  |
| P value | <0.0001 |  |  |  |  |
| P value summary | **** |  |  |  |  |
| Significant diff. among means (P < 0.05)? | Yes |  |  |  |  |
| Tukey's multiple comparisons test | Mean diff. | | 95.00% CI of diff. | Adjusted *P* Value | |
| WT vs. KO HET | -114.4 | | -171.1 to -57.74 | <0.0001 | |
| WT vs. G34S HET | -2.387 | | -56.22 to 51.45 | 0.9939 | |
| KO HET vs. G34S HET | 112.0 | | 53.07 to 171.0 | <0.0001 | |
| **Male** |  | | | | |
| ANOVA summary |  |  | | | |
| F | 9.840 |  |  |  |  |
| P value | 0.0001 |  |  |  |  |
| P value summary | *** |  |  |  |  |
| Significant diff. among means (P < 0.05)? | Yes |  |  |  |  |
| Tukey's multiple comparisons test | Mean diff. | | 95.00% CI of diff. | | Adjusted *P* Value |
| WT vs. KO HET | -85.90 | | -135.6 to -36.17 | | 0.0003 |
| WT vs. G34S HET | 2.320 | | -45.38 to 50.02 | | 0.9926 |
| KO HET vs. G34S HET | 88.22 | | 32.55 to 143.9 | | 0.0008 |

**Supplementary Table 5. Statistical analysis of the time spent inside.**

| **Female** |  | | | | |
| --- | --- | --- | --- | --- | --- |
| ANOVA summary |  |  | | | |
| F | 14.23 |  |  |  |  |
| P value | <0.0001 |  |  |  |  |
| P value summary | **** |  |  |  |  |
| Significant diff. among means (P < 0.05)? | Yes |  |  |  |  |
| Tukey's multiple comparisons test | Mean diff. | | 95.00% CI of diff. | Adjusted *P* Value | |
| WT vs. KO HET | 112.2 | | 56.41 to 168.0 | <0.0001 | |
| WT vs. G34S HET | -1.259 | | -54.83 to 52.31 | 0.9983 | |
| KO HET vs. G34S HET | -113.5 | | -171.9 to -55.04 | <0.0001 | |
| **Male** |  | | | | |
| ANOVA summary |  |  | | | |
| F | 9.840 |  |  |  |  |
| P value | 0.0001 |  |  |  |  |
| P value summary | *** |  |  |  |  |
| Significant diff. among means (P < 0.05)? | Yes |  |  |  |  |
| Tukey's multiple comparisons test | Mean diff. | | 95.00% CI of diff. | | Adjusted *P* Value |
| WT vs. KO HET | 85.90 | | 36.17 to 135.6 | | 0.0003 |
| WT vs. G34S HET | -2.322 | | -50.02 to 45.38 | | 0.9926 |
| KO HET vs. G34S HET | -88.22 | | -143.9 to -32.55 | | 0.0008 |

**Supplementary Table 6. Statistical analysis of contextual fear conditioning.**

| **Female** |  | | | | |
| --- | --- | --- | --- | --- | --- |
| ANOVA summary |  |  | | | |
| F | 4.843 |  |  |  |  |
| P value | 0.0141 |  |  |  |  |
| P value summary | * |  |  |  |  |
| Significant diff. among means (P < 0.05)? | Yes |  |  |  |  |
| Tukey's multiple comparisons test | Mean diff. | | 95.00% CI of diff. | Adjusted *P* Value | |
| WT vs. KO HET | 12.34 | | 1.622 to 23.06 | 0.0211 | |
| WT vs. G34S HET | 11.92 | | 0.2166 to 23.62 | 0.0452 | |
| KO HET vs. G34S HET | -0.4185 | | -11.94 to 11.10 | 0.9956 | |
| **Male** |  | | | | |
| ANOVA summary |  |  | | | |
| F | 8.184 |  |  |  |  |
| P value | 0.0013 |  |  |  |  |
| P value summary | ** |  |  |  |  |
| Significant diff. among means (P < 0.05)? | Yes |  |  |  |  |
| Tukey's multiple comparisons test | Mean diff. | | 95.00% CI of diff. | | Adjusted *P* Value |
| WT vs. KO HET | 13.51 | | 2.105 to 24.92 | | 0.0174 |
| WT vs. G34S HET | 16.58 | | 5.543 to 27.62 | | 0.0023 |
| KO HET vs. G34S HET | 3.067 | | -9.515 to 15.65 | | 0.8216 |

**Supplementary Table 7. Statistical analysis of immunoblot analysis in Fig. 5a and 5b.**

| **Female δ-catenin** |  | | | | |
| --- | --- | --- | --- | --- | --- |
| ANOVA summary |  |  | | | |
| F | 10.63 |  |  |  |  |
| P value | 0.0003 |  |  |  |  |
| P value summary | *** |  |  |  |  |
| Significant diff. among means (P < 0.05)? | Yes |  |  |  |  |
| Tukey's multiple comparisons test | Mean diff. | | 95.00% CI of diff. | Adjusted *P* Value | |
| WT vs. KO HET | 0.5074 | | 0.1364 to 0.8784 | 0.0055 | |
| WT vs. G34S HET | -0.1602 | | -0.5312 to 0.2108 | 0.5453 | |
| KO HET vs. G34S HET | -0.6676 | | -1.039 to -0.2966 | 0.0003 | |
| **Male δ-catenin** |  | | | | |
| ANOVA summary |  |  | | | |
| F | 4.211 |  |  |  |  |
| P value | 0.0235 |  |  |  |  |
| P value summary | * |  |  |  |  |
| Significant diff. among means (P < 0.05)? | Yes |  |  |  |  |
| Tukey's multiple comparisons test | Mean diff. | | 95.00% CI of diff. | | Adjusted *P* Value |
| WT vs. KO HET | 0.3152 | | 0.01229 to 0.6181 | | 0.0400 |
| WT vs. G34S HET | 0.01017 | | -0.2927 to 0.3131 | | 0.9963 |
| KO HET vs. G34S HET | -0.3050 | | -0.6080 to -0.002116 | | 0.0481 |
| **Female GluA1** |  | |  | |  |
| ANOVA summary |  | |  | |  |
| F | 1.425 | |  | |  |
| P value | 0.2550 | |  | |  |
| P value summary | ns | |  | |  |
| Significant diff. among means (P < 0.05)? | No | |  | |  |
| Tukey's multiple comparisons test | Mean diff. | | 95.00% CI of diff. | | Adjusted *P* Value |
| WT vs. KO HET | 0.2700 | | -0.1745 to 0.7145 | | 0.3085 |
| WT vs. G34S HET | 0.2593 | | -0.1852 to 0.7038 | | 0.3368 |
| KO HET vs. G34S HET | -0.01066 | | -0.4552 to 0.4339 | | 0.9981 |
| **Male GluA1** |  | |  | |  |
| ANOVA summary |  | |  | |  |
| F | 0.7501 | |  | |  |
| P value | 0.4802 | |  | |  |
| P value summary | ns | |  | |  |
| Significant diff. among means (P < 0.05)? | No | |  | |  |
| Tukey's multiple comparisons test | Mean diff. | | 95.00% CI of diff. | | Adjusted *P* Value |
| WT vs. KO HET | 0.08619 | | -0.3650 to 0.5374 | | 0.8864 |
| WT vs. G34S HET | -0.1371 | | -0.5883 to 0.3141 | | 0.7384 |
| KO HET vs. G34S HET | -0.2233 | | -0.6745 to 0.2279 | | 0.4533 |
| **Female GluA2** |  | |  | |  |
| ANOVA summary |  | |  | |  |
| F | 0.7975 | |  | |  |
| P value | 0.4589 | |  | |  |
| P value summary | ns | |  | |  |
| Significant diff. among means (P < 0.05)? | No | |  | |  |
| Tukey's multiple comparisons test | Mean diff. | | 95.00% CI of diff. | | Adjusted *P* Value |
| WT vs. KO HET | 0.1050 | | -0.5309 to 0.7410 | | 0.9137 |
| WT vs. G34S HET | -0.2160 | | -0.8519 to 0.4200 | | 0.6853 |
| KO HET vs. G34S HET | -0.3210 | | -0.9569 to 0.3150 | | 0.4395 |
| **Male GluA2** |  | |  | |  |
| ANOVA summary |  | |  | |  |
| F | 0.8480 | |  | |  |
| P value | 0.4374 | |  | |  |
| P value summary | ns | |  | |  |
| Significant diff. among means (P < 0.05)? | No | |  | |  |
| Tukey's multiple comparisons test | Mean diff. | | 95.00% CI of diff. | | Adjusted *P* Value |
| WT vs. KO HET | -0.1539 | | -0.5590 to 0.2511 | | 0.6239 |
| WT vs. G34S HET | -0.2069 | | -0.6120 to 0.1981 | | 0.4310 |
| KO HET vs. G34S HET | -0.05299 | | -0.4581 to 0.3521 | | 0.9449 |

**Supplementary Table 8. Statistical analysis of immunoblot analysis in Fig. 5c and 5d.**

| **Female δ-catenin** |  | | | | |
| --- | --- | --- | --- | --- | --- |
| ANOVA summary |  |  | | | |
| F | 17.21 |  |  |  |  |
| P value | <0.0001 |  |  |  |  |
| P value summary | **** |  |  |  |  |
| Significant diff. among means (P < 0.05)? | Yes |  |  |  |  |
| Tukey's multiple comparisons test | Mean diff. | | 95.00% CI of diff. | Adjusted *P* Value | |
| WT vs. KO HET | 0.6197 | | 0.3529 to 0.8866 | <0.0001 | |
| WT vs. G34S HET | 0.1781 | | -0.08874 to 0.4450 | 0.2443 | |
| KO HET vs. G34S HET | -0.4416 | | -0.7085 to -0.1747 | 0.0008 | |
| **Male δ-catenin** |  | | | | |
| ANOVA summary |  |  | | | |
| F | 5.072 |  |  |  |  |
| P value | 0.0120 |  |  |  |  |
| P value summary | * |  |  |  |  |
| Significant diff. among means (P < 0.05)? | Yes |  |  |  |  |
| Tukey's multiple comparisons test | Mean diff. | | 95.00% CI of diff. | | Adjusted *P* Value |
| WT vs. KO HET | 0.3201 | | 0.03208 to 0.6082 | | 0.0267 |
| WT vs. G34S HET | 0.3273 | | 0.03926 to 0.6154 | | 0.0231 |
| KO HET vs. G34S HET | 0.007176 | | -0.2809 to 0.2952 | | 0.9979 |
| **Female GluA1** |  | |  | |  |
| ANOVA summary |  | |  | |  |
| F | 6.959 | |  | |  |
| P value | 0.0030 | |  | |  |
| P value summary | ** | |  | |  |
| Significant diff. among means (P < 0.05)? | Yes | |  | |  |
| Tukey's multiple comparisons test | Mean diff. | | 95.00% CI of diff. | | Adjusted *P* Value |
| WT vs. KO HET | 0.1805 | | -0.3684 to 0.7294 | | 0.7014 |
| WT vs. G34S HET | -0.6154 | | -1.164 to -0.06647 | | 0.0253 |
| KO HET vs. G34S HET | -0.7959 | | -1.345 to -0.2470 | | 0.0032 |
| **Male GluA1** |  | |  | |  |
| ANOVA summary |  | |  | |  |
| F | 7.420 | |  | |  |
| P value | 0.0022 | |  | |  |
| P value summary | ** | |  | |  |
| Significant diff. among means (P < 0.05)? | Yes | |  | |  |
| Tukey's multiple comparisons test | Mean diff. | | 95.00% CI of diff. | | Adjusted *P* Value |
| WT vs. KO HET | 0.2464 | | -0.2779 to 0.7707 | | 0.4890 |
| WT vs. G34S HET | -0.5569 | | -1.081 to -0.03262 | | 0.0354 |
| KO HET vs. G34S HET | -0.8033 | | -1.328 to -0.2790 | | 0.0019 |
| **Female GluA2** |  | |  | |  |
| ANOVA summary |  | |  | |  |
| F | 6.282 | |  | |  |
| P value | 0.0049 | |  | |  |
| P value summary | ** | |  | |  |
| Significant diff. among means (P < 0.05)? | Yes | |  | |  |
| Tukey's multiple comparisons test | Mean diff. | | 95.00% CI of diff. | | Adjusted *P* Value |
| WT vs. KO HET | 0.3291 | | 0.07623 to 0.5821 | | 0.0084 |
| WT vs. G34S HET | 0.02725 | | -0.2257 to 0.2802 | | 0.9623 |
| KO HET vs. G34S HET | -0.3019 | | -0.5548 to -0.04898 | | 0.0164 |
| **Male GluA2** |  | |  | |  |
| ANOVA summary |  | |  | |  |
| F | 8.514 | |  | |  |
| P value | 0.0010 | |  | |  |
| P value summary | ** | |  | |  |
| Significant diff. among means (P < 0.05)? | Yes | |  | |  |
| Tukey's multiple comparisons test | Mean diff. | | 95.00% CI of diff. | | Adjusted *P* Value |
| WT vs. KO HET | 0.3066 | | 0.01420 to 0.5989 | | 0.0382 |
| WT vs. G34S HET | -0.1796 | | -0.4719 to 0.1128 | | 0.3006 |
| KO HET vs. G34S HET | -0.4861 | | -0.7785 to -0.1938 | | 0.0008 |
